# Characterization of aroma-active compounds in delicious apples juices by gas chromatography-mass spectrometry (GC-MS), gas chromatography–olfactometry (GC-O) and odor activity value (OAV)

**DOI:** 10.1101/611061

**Authors:** Mao deshou, Hong liu, Li zhengfeng, Niu yunwei, Xiao zuobing, Zhang fengmei, Zhu jiancai

## Abstract

Volatile aroma compounds of delicious apple juice in three cultivars were obtained by gas chromatography-mass spectrometry (GC-MS), gas chromatography-olfactometry (GC-O), and GC-flame photometric detection (FPD). Quantitatively, the major volatiles of the delicious apple juice were detected by GC-MS, such as esters, alcohols, aldehydes. In addition, GC-O and OAV were used to determine the aroma-active compounds in fruit. Amongst these compounds, ethyl 2-methylbutanoate (47-229), butyl 2-methylbutanoate (8-208), (E)-2-hexenal (25-120), butyl propanoate (14-54), methyl 2-methylbutanoate (28-41), ethyl hexanoate (4-32), ethyl butanoate (5-17) showed high OAVs in three delicious apple juices, which contributed greatly to the aroma of delicious apple juice. Beside those compounds, methanethiol (OAV: 1.1-1.6), dimethyl sulfide (OAV: 2.5-3.6), methional (OAV: 4.2-11.7) and 2-(methylthio)ethanol (OAV: 1.2-1.9) also presented relatively high OAVs. Finally, four compounds (ethyl 2-methylbutanoate, ethyl octanoate, ethyl butanoate and ethyl hexanoate) were selected to investigate the possible interactions occurring in the delicious apple juice. The resultants demonstrated that those aroma volatile compounds can decrease threshold of the solution to dissimilar degrees before and after their addition.

## 1. Introduction

Aroma was used as significant index for determination the quality in natural products. There was a variety of volatile compounds in natural products including esters, terpenols, alcohols, *etc.* With different aroma features and contents, these compounds formed the aroma of natural products. Gas chromatography (GC) in conjunction with mass selective detector (MSD) were widely employed to identify volatile compounds in natural products, such as mango (Pino & Mesa., 2010), pink guava (Martin et al., 2009), orange(Kanjana et al., 2005).

In addition to these compounds, sulfur compounds might also contribute significantly to the aromas of fruits. For example, dimethyl disulfide, 3-mercaptohexanol and 4-mercapto-4-methyl-2-pentanone were great contributors in cranberry fruits (Zhu et al., 2016). In addition, 2-isopropyl-4-methylthiazole and 4-mercapto-4-methyl-2-pentanone played important role in aroma of peach (Zhu & Xiao., 2019). 3-Mercapto-3-methylbut-1-ol and furfuryl mercaptan were also characteristic compounds in raspberries (Duarte et al., 2010). Although the content of sulfur in fruit was very low, the contribution of those compounds to the aroma of fruits was significant because of the extremely low odor threshold.

The delicious apple was rich in reducing sugar, protein, phosphorus, iron, malic acid, quinic acid and citric acid. Recently, many researches focused on the aroma of delicious apple (Fallik et al., 1997; Song & Bangerth, 1996; Mattheis et al., 1991; Rowan et al., 1999; Streif & Bangerth., 1988; Mattheis et al., 1995). For example, Fallik et al studied the influence of heat treatment to inhibit compounds emission in delicious apples (Fallik et al., 1997). Volatile compounds released from delicious apple during transition period were also identified (Mattheis et al., 1991). Although some research works had been done on delicious apples, previous researches just focused on the quantitation of compounds in delicious apples. The key aroma compounds in delicious apples were not identified. Therefore, the purposes of this investigation were to identify volatile compounds by GC-MS and FPD in delicious apples, and to determine the aroma-active compounds with GC-O and OAV, and finally, to investigate interactions between different aroma compounds in delicious apples.

## 2 Materials and methods

### 2.1 Chemicals

The standards compounds were purchased from Aladdin Corporation (Tianjin, China). The n-alkanes (C_6_-C_30_) standard were obtained from Sigma-Aldrich (Shanghai, China). All the standards compounds were of GC quality.

### 2.2 Preparation of sample

Three red delicious apple cultivars (*Malus domestica*) harvested from different areas were studied. Sample Y1 was purchased in Wal-Mart which imported from State of Washington (USA). Sample Y2 and Y3 were obtained from local farm in Shangdong and Shan’Xi Province (China), which were harvested in 2017. The fruits were selected with red color. Specifically, 1kg of fruits were deseeded. Then, the samples and deionised water (500 g) were mixed together in blender. After 5 minutes, the mixtures were prepared. The mixtures were filtered to get the supernatant. Last, delicious apple juices were saved in a refrigerator with 4 ° C.

### 2.3 Soild phase micro-extration (SPME) absorption of volatile compounds in delicious apple juices

The volatile compounds in delicious apple juices were analyzed by SPME. The type of fiber was 75 μm carboxyl/poly dimethyl siloxane (CAR/PDMS). Then, 8 mL of the treated samples and 0.01 mL of 2-octanol (5 mg/L) were mixed in vials and covered. Then, the vials were placed in the water bath with 30 °C for 5 minutes. After that, the fibre was put on top of the headspace vial for 30 min. Then, the fibre was withdrawn and inserted in the injector of 7890-5975 GC-MS. All experiments were repeated three times.

### 2.4 Preparation of model solution

The prepared delicious apple juices were gradually flushed with ultra-high purity (99.99%) nitrogen to eliminate volatile compounds until none volatile compound was detected with GC-MS. The nitrogen rate was set at 1 mL/min. Finally, the model solution for standard curves was prepared.

### 2.5 Calibration of standard curves

In this process, stock solution consisted of compounds in delicious apple juice were prepared. The stock solution was diluted to 5, 10, 20, 30, 40 and 50 gradients with water. Then, 2-octanol and dipropyl disulfide were used as standard compounds for quantitation the non-sulfur and sulfur compounds, respectively. Six diluted solutions (0.01 g) and internal standard solution (0.01 g) with 2-octanol (5 mg/kg) or dipropyl disulfide (10 μg/kg) were placed in vial with 8 mL of model solution. Finally, the model solution was analysed with SPME. The aroma compounds were quantified by external standard curves. All experiments were repeated three times.

### 2.6 GC-MS of volatile compounds in delicious apple juices

GC-MS (7890-5975) was employed to analyze volatile compounds of delicious apple juices. Two different analytical fused silica capillary columns, HP-Innowax and DB-5, were employed in the experiment with the type of 60 m (length)× 0.25 mm (diam)×0.25 μm (film). The temperature of injection was 250 °C. The rate of carrier gas (Helium) was 1 mL/min. The electron impact energy, ion source temperature and quadrupole mass filter were 70 eV, 230 °C and 150 °C, respectively. The initial temperature of oven was 40 °C and kept for 6 min, then programed to 100 °C (3 °C/ min) and programed to 230 °C (5 °C/ min). Finally, the 230 °C was hold for 10 min. The compounds in the delicious apple juices were identified with standard compounds, retention indices (RIs) and mass spectra in the Database (Wiley7n.l). The RIs were calculated by alkane mixtures (C_6_-C_30_).

### 2.7 Descriptive testing methods

The sensory of delicious apple juices were determined by panel of 10 members from sensory laboratory with ages of 25-45. The detailed information of sensory training was referred to previous literature (Zhu et al., 2018). There were few research about the sensory profile of delicious apple. Thus, the identification of sensory descriptors of delicious apple juices was referred to the apple due to the similar aroma between apple and delicious apple (Seppä et al., 2012). Furthermore, “Aroma Wheel” was also employed to identify the sensory descriptors of delicious apple. The characteristic notes of delicious apple was determined as six descriptors including wax, sweet, fruit, green, floral and sour. Those descriptors were defined as standard compounds. Hexanol represented for “wax” descriptor, maltol for “sweet” descriptor, (E)-2-hexenal for “green” descriptor, ethyl hexanoate for “fruit” descriptor, phenylethyl alcohol for “floral” descriptor, acetic acid for “sour” descriptor, dipropyl disulfide for “sulfur” descriptor. The scale of 10 points was used to evaluate the intensity of descriptors. “0” referred to undetected, “3” referred to weak, “5” referred to moderate, “10” referred to strong. All experiments were repeated three times.

### 2.8 SPME-GC-FID-O analysis of delicious apple juices

GC-O analysis was employed on Agilent (7890) GC equipped with an olfactometer (ODP3, Gerstel, Germany). The information of columns and oven program were identical to the conditions of GC-MS. The effluent flowed into FID and the heated olfactory at ratio of 1:1. Ten trained panelists (5 males and 5 females) participated in GC-O experiment. As described in Section 2.5, those panelists has been trained with standard compounds. And the odor intensities were also evaluated using 10-point intensity scale. When the panelists sniffed the mask effluent, they recorded the start and end times of the effluent of the aromatic active compound, as well as the characteristics and intensity of the odor. The experiment was repeated three times, and the final intensity of the aroma compound was averaged among ten panelists.

### 2.9 SPME-GC-FPD

Flame photometric detector (FPD) was employed to detect sulfur compounds in delicious apple juices. The columns and oven program were identical to the GC-MS. The temperature of FPD was 250 °C. Those sulfur compounds were determined on the basis of authentic standards and retention index.

### 2.10 Odor activity value (OAV)

The OAV was employed to evaluate contribution of compounds to aroma of delicious apple juices. The threshold values was referred to available reference (Van., 2003).

### 2.11 The comparing of detection probability of reconstitution solution before and after adding volatile compounds

The sensory panel measured the thresholds of reconstitution solutions before, and after, omitting certain compounds using three alternative, forced-choice presentation (3-AFC). The complete reconstitution solution consisted of volatile compounds in delicious apple juices of Y1 sample with actual concentration. To begin with, four compounds (ethyl 2-methylbutanoate, ethyl hexanoate, ethyl butanoate, and ethyl octanoate) were omitted from complete solution separately. The volume of reconstitution solutions replaced the concentration. And initial reconstitution solution volume was 0.1mL. The solution was diluted with water in multiples of 2. Ten volume gradients were used. In this experiment, sigmoid curve was employed to determine detection threshold (Lytra et al., 2013). Detection threshold is defined as the corresponding volume when the correct detection probability was 50%. The change of sensory intensity was also evaluated before and after the addition of ethyl 2-methylbutanoate, ethyl hexanoate, ethyl butanoate, and ethyl octanoate.

### 2.12 Statistical analysis

Variance analysis (ANOVA) was employed to judge the concentrations and descriptors differences between samples. Origin 8.0 software was used for sigmoid curve.

## 3 Results and conclusions

### 3.1 Optimized HS-SPME conditions

Before extraction, several parametres in SPME should be investigated, such as sample amount, extraction time and extraction temperature. The amount of volatile compounds absorbed on SPME fibers might be related to the amount of samples. Normally, the larger the sample amount, the more volatile compounds absorbed by the extraction fiber. However, too many samples could weaken volatilization of volatile compounds. At the same time, the adsorption of fibers tends to be saturated, which was not conducive to the extraction of volatile compounds (Rocha et al. 2001). In this experiment, sample amounts of 3, 4, 5 and 6 g were investigated. The results showed that the optimum extraction conditions was 5 g.

Extraction time was an another important parameter, which could affect extraction efficiency in experiment. If the extraction time of the fibers in the headspace bottle was too long, the competition between the volatile compounds on the limited position of the fibers would lead to the inaccuracy of the relative content of the analytes (Howard et al., 2005). In this experiment, the extraction time was 15, 30, 45 and 60 minutes, respectively. The results demonstrated that the optimum extraction time was 45 minutes. Extraction temperature was also an important factor in SPME experiment. Volatile compounds in fruits are particularly sensitive to temperature. It is easy to produce various by-products during high temperature, especially sulfur compounds (Chung-May & Wang., 2000). Therefore, the temperature of 25, 30 and 35 °C were investigated. The results suggested that the volatile compounds increased with the extraction temperature elevated. After comprehensive consideration, the extraction temperature should be 30 °C. Based on these observations, the optimum conditions of HS-SPME were determined, i.e. extraction time 45 minutes, sample volume 5 g and extraction temperature 30 °C.

### 3.2 GC-O results for delicious apple juices

The results of the GC-O analysis were listed in Table 1. Application of GC-O on the delicious apple juices revealed 40, 37 and 32 volatile compounds in Y1, Y2 and Y3 samples, respectively. The different aroma intensities (AIs) of the volatile compounds in each of the samples were mainly caused by concentration differences of these compounds. The AIs of the compounds ranged from 0.1 to 8.5 for Y1, 0.1 to 8.6 for Y2, 0.1 to 7.3 for Y3, respectively. From the Table 1, (E)-2-hexenal was present at the highest AI in Y1 sample. Ethyl 2-methylbutanoate presented the highest AIs in the Y2 (AI=8.6) and Y3 (AI=7.3) samples.

**Table 1.** GC-O identified aroma-active compounds in three delicious apple juices with the method of aroma intensity.

| No | Compounds<sup>A</sup> | RI<sup>B</sup> |  | Identification<sup>C</sup> | Aroma description | Aroma intensity |  |  |  |  |  |
| --- | --- | --- | --- | --- | --- | --- | --- | --- | --- | --- | --- |
|  |  | Innowax | DB-5 |  |  | Y1 | RSD(%) | Y2 | RSD(%) | Y3 | RSD(%) |
| 1 | Methanethiol | 694 | 504 | AD, RI, Std | sulfur, gasoline | 2.3 | 4.5 | 2.6 | 5.4 | 1.9 | 3.6 |
| 2 | Dimethyl sulfide | 717 | 522 | AD, RI, Std | sulfur, corn | 3.8 | 6.5 | 3.2 | 3.6 | 4.3 | 5.4 |
| 3 | Ethanethiol | 720 | 512 | AD, RI, Std | sulfur, garlic | 0.6 | 4.6 | 0.8 | 3.2 | 0.7 | 4.6 |
| 4 | Butanal | 833 | 599 | AD, RI, Std | apple | 3.5 | 6.5 | 1.9 | 6.1 | - | - |
| 5 | Ethyl acetate | 909 | 629 | AD, RI, Std | paint, fruity | 0.6 | 4.7 | 0.4 | 4.6 | 0.1 | 4.8 |
| 6 | Pentanal | 938 | 734 | AD, RI, Std | almond, malt | 1.2 | 5.4 | - | - | - | - |
| 7 | Ethyl propanoate | 950 | 712 | AD, RI, Std | fruity | 4.1 | 6.7 | 3.9 | 6.3 | 6.8 | 7.0 |
| 8 | Propyl acetate | 955 | 652 | AD, RI, Std | fruity | 0.3 | 4.8 | 0.5 | 4.3 | - | - |
| 9 | Ethyl isobutanoate | 957 | 758 | AD, RI, Std | fruity, apple | 4.8 | 7.1 | 5.7 | 7.4 | 6.4 | 6.3 |
| 10 | Methyl butanoate | 992 | 726 | AD, RI, Std | fruity, sweet | -<sup>D</sup> | - | 5.6 | 3.9 | 4.7 | 4.7 |
| 11 | Methyl 2-methylbutanoate | 1008 | 738 | AD, RI, Std | fruity | 7.2 | 5.6 | 7.5 | 5.8 | 7.1 | 6.2 |
| 12 | Unknown1 | 1016 | 777 | AD | floral | - | - | 0.2 | 6.5 | - | - |
| 13 | Butanol | 1026 | 674 | AD, RI, Std | green, wine | 0.2 | 4.2 | 0.1 | 4.5 | 0.2 | 4.4 |
| 14 | Ethyl butanoate | 1027 | 806 | AD, RI, Std | fruity, apple | 5.8 | 5.1 | 6.3 | 5.3 | 3.7 | 4.7 |
| 15 | Unknown2 | 1029 | 627 | AD | green | 0.1 | 6.5 | - | - | - | - |
| 16 | Unknown3 | 1043 | 699 | AD | sweet | 2.3 | 4.3 | 3.5 | 3.7 | - | - |
| 17 | Ethyl 2-methylbutanoate | 1061 | 845 | AD, RI, Std | sweet caramel, grape | 7.8 | 4.5 | 8.6 | 4.1 | 7.3 | 4.0 |
| 18 | Butyl acetate | 1074 | 815 | AD, RI, Std | fruity, apple | 5.7 | 4.8 | 6.7 | 4.3 | 5.5 | 4.5 |
| 19 | Hexanal | 1079 | 805 | AD, RI, Std | grass, tallow, fat | 4.7 | 5.0 | 4.8 | 4.7 | 4.3 | 4.8 |
| 20 | 2-Methylpropanol | 1097 | 650 | AD, RI, Std | green | 0.2 | 4.9 | - | 4.8 | 0.4 | 5.0 |
| 21 | Unknown4 | 1133 | 755 | AD | leaf, green | 2.1 | 3.4 | 2.3 | 4.3 | 1.9 | 2.9 |
| 22 | Ethyl pentanoate | 1136 | 905 | AD, RI, Std | fruity | 0.1 | 5.6 | - | - | - | - |
| 23 | Butyl propanoate | 1139 | 907 | AD, RI, Std | fruity | 6.5 | 7.0 | 6.3 | 7.3 | 5.6 | 6.2 |
| 24 | (E)-2-Hexenal | 1194 | 842 | AD, RI, Std | green, leaf | 8.5 | 7.5 | 6.4 | 6.7 | 6.6 | 6.3 |
| 25 | 2-Methylbutanol | 1207 | 670 | AD, RI, Std | mellow,herbaceous | 0.3 | 4.2 | 0.5 | 3.5 | 0.4 | 4.3 |
| 26 | Ethyl hexanoate | 1222 | 1005 | AD, RI, Std | apple peel, fruity | 5.5 | 5.2 | 6.8 | 5.3 | 4.5 | 5.7 |
| 27 | Butyl 2-methylbutanoate | 1223 | 1042 | AD, RI, Std | fruity, wine | 5.4 | 4.5 | 7.4 | 5.0 | 3.6 | 4.0 |
| 28 | Butyl butanoate | 1223 | 991 | AD, RI, Std | fruity, wine | 4.5 | 4.4 | 5.4 | 3.9 | 3.2 | 3.9 |
| 29 | (E)-2-Heptenal | 1244 | 955 | AD, RI, Std | soap, fat | 0.3 | 4.0 | - | - | - | - |
| 30 | Heptanol | 1271 | 928 | AD, RI, Std | green,vegetal | 0.4 | 5.5 | 0.3 | 5.1 | - | - |
| 31 | Hexanol | 1362 | 852 | AD, RI, Std | vegetal, herbaceous | 0.4 | 7.2 | - | - | 3.4 | 7.3 |
| 32 | (E)-2-Hexen-1-ol | 1380 | 855 | AD, RI, Std | leaf, green | - | - | 0.3 | 7.2 | 0.4 | 6.7 |
| 33 | Nonanal | 1388 | 1105 | AD, RI, Std | fat, citrus, green | 4.3 | 4.3 | 4.2 | 4.1 | 0.2 | 4.5 |
| 34 | Octanol | 1390 | 979 | AD, RI, Std | critus, fat | 3.7 | 4.6 | 3.4 | 4.8 | - | - |
| 35 | Unknown5 | 1394 | 860 | AD | fat | 0.3 | 3.8 | - | - | - | - |
| 36 | Methional | 1446 | 907 | AD, RI, Std | cooked potato | 3.4 | 4.4 | 4.5 | 3.7 | 5.1 | 5.1 |
| 37 | Unknown6 | 1449 | 603 | AD | critus | - | - | 1.7 | 4.3 | 2.1 | 4.6 |
| 38 | Acetic acid | 1450 | 600 | AD | acid | 1.9 | 3.6 | 2.1 | 4.3 | 2.4 | 5.5 |
| 39 | Benzaldehyde | 1499 | 961 | AD, RI, Std | almond, burnt sugar | 4.5 | 4.5 | 3.2 | 4.6 | 2.8 | 5.0 |
| 40 | Nonanol | 1500 | 1155 | AD, RI, Std | fat, green | 0.4 | 4.9 | 0.6 | 5.5 | - | - |
| 41 | Benzyl acetate | 1512 | 1160 | AD, RI, Std | floral | 6.5 | 6.4 | 7.1 | 6.9 | 5.7 | 6.6 |
| 42 | 2-(Methylthio)ethanol | 1515 | 820 | AD, RI, Std | sulfur | 1.2 | 4.6 | 1.5 | 5.4 | 0.6 | 4.5 |
| 43 | Methionol | 1722 | 987 | AD, RI, Std | vegetable, potato | 0.3 | 2.7 | 0.1 | 2.9 | 0.2 | 3.2 |
| 44 | β-Damascenone | 1818 | 1389 | AD, RI, Std | $\beta$ -Damascenone | 5.5 | 6.7 | 5.2 | 6.9 | 4.6 | 6.1 |

<sup>A</sup> Volatile compounds detected in the delicious apple juices samples; <sup>B</sup> Retention index of compounds on DB-5 and Innowax Column; <sup>C</sup> RI: retention index; Std:
confirmed by authentic standards; AD: Aroma descriptor; <sup>D</sup> -: not perceive;

Ester compounds were shown to be the largest class of aroma compounds in the delicious apple juices. A total of 14 esters were identified, which were summarized in Table 1. These included ethyl butanoate, ethyl 2-methylbutanoate, methyl 2-methylbutanoate, butyl propanoate, ethyl isobutanoate, ethyl hexanoate, butyl 2-methylbutanoate, and butyl butanoate. As is well-known, ester compounds were considered as fruity, apple notes in the GC-O analysis (Table 1). There was no doubted that these esters form the fundamental of the aroma of the fruit. Amongst esters, ethyl 2-methylbutanoate, methyl 2-methylbutanoate, butyl propanoate and ethyl butanoate were the most powerful aroma-active compounds, which contributed to the aroma of delicious apple juices. The results were agreed with previous studies, which demonstrated esters contributed greatly to fruits with characteristic fruity aroma (Du & Rouseff., 2014; Pang et al., 2012).

In addition to those ester compounds, aldehydes were another important class of aroma active compounds in the delicious apple juices. Amongst those compounds, (E)-2-hexenal, hexanal and nonanal contributed greatly to the aroma profiles of delicious apple juices. According to GC-O analysis, these compounds were considered as fatty, green, leaf notes, which provided fresh profile to the fruits.

Beside those non-sulfur compounds, sulfur volatile compounds also perceived in experiment, such as methional, ethanethiol, methanethiol, dimethyl sulfide, 2-(methylthio)ethanol and methionol. Amongst those sulfur compounds, methional, ethanethiol, dimethyl sulfide presented high AIs, which considered as important compounds to delicious apple juices.

### 3.3 Quantitative analysis and OAV of volatile compounds

#### 3.3.1 Quantitative analysis and OAV of sulfur compounds

As listed in Table 2, a total of nine sulfur compounds were identified in two dissimilar columns by FPD. The calibration curves, determination coefficients, validation range, LOD (μg/kg) and LOQ (μg/kg) were established in this experiment. The results demonstrated that those indicator were all valid. From the Table 3, 8, 9, and 9 sulfur compounds were detected in three samples, respectively. Quantitatively, 2-(methylthio)ethanol, methionol, methanethiol, 3-methyl-2-butene-1-thiol presented relatively higher contents than other sulfur compounds. However, ethanethiol, propanethiol and methional presented relatively low contents in delicious apple juices. According to the previous investigations, the contributions of compounds to global aroma was not determined by content, but by OAV of compounds. Normally, the compounds were considered as contributors to aroma with OAV above 1 (Guth, 1997). From Table 3, methanethiol (OAV: 1.1-1.6), dimethyl sulfide (OAV: 2.5-3.6), methional (OAV: 4.2-11.7) and 2-(methylthio)ethanol (OAV: 1.2-1.9) presented relatively high OAVs.

**Table 2.**
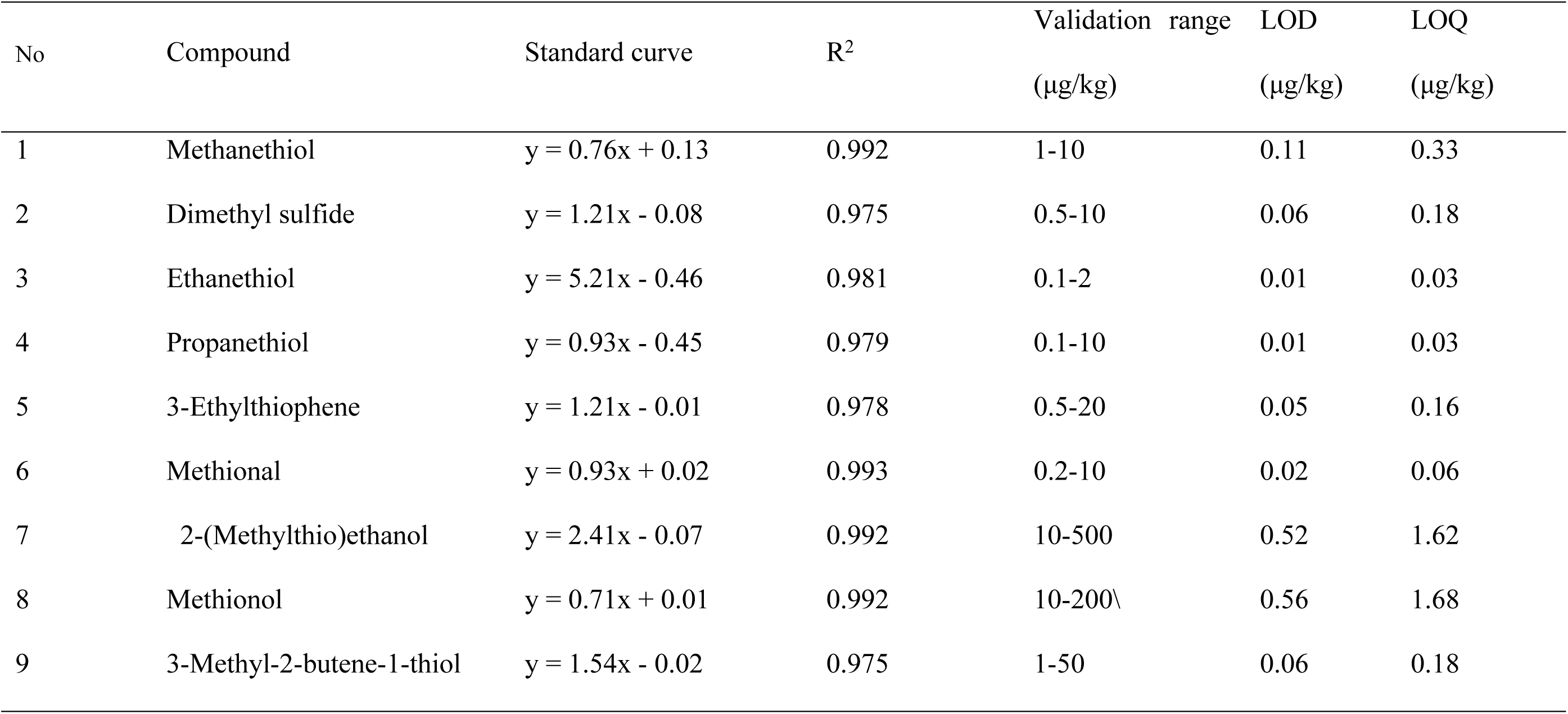
The standard curve, validation range and determination coefficients (R^2^) for the volatile compounds in three delicious apple juices.

| No | Compound | Standard curve | $R^2$ | Validation range<br>($\mu\text{g/kg}$) | LOD<br>($\mu\text{g/kg}$) | LOQ<br>($\mu\text{g/kg}$) |
| --- | --- | --- | --- | --- | --- | --- |
| 1 | Methanethiol | $y = 0.76x + 0.13$ | 0.992 | 1-10 | 0.11 | 0.33 |
| 2 | Dimethyl sulfide | $y = 1.21x - 0.08$ | 0.975 | 0.5-10 | 0.06 | 0.18 |
| 3 | Ethanethiol | $y = 5.21x - 0.46$ | 0.981 | 0.1-2 | 0.01 | 0.03 |
| 4 | Propanethiol | $y = 0.93x - 0.45$ | 0.979 | 0.1-10 | 0.01 | 0.03 |
| 5 | 3-Ethylthiophene | $y = 1.21x - 0.01$ | 0.978 | 0.5-20 | 0.05 | 0.16 |
| 6 | Methional | $y = 0.93x + 0.02$ | 0.993 | 0.2-10 | 0.02 | 0.06 |
| 7 | 2-(Methylthio)ethanol | $y = 2.41x - 0.07$ | 0.992 | 10-500 | 0.52 | 1.62 |
| 8 | Methionol | $y = 0.71x + 0.01$ | 0.992 | 10-200\ | 0.56 | 1.68 |
| 9 | 3-Methyl-2-butene-1-thiol | $y = 1.54x - 0.02$ | 0.975 | 1-50 | 0.06 | 0.18 |

**Table 3.** The concentration (mg/kg), standard deviation and OAV of aroma compounds detected in three delicious apple juices

| No | Compound | RI<sup>A</sup> |  | Identification<sup>B</sup> | Y1 |  | Y2 |  | Y3 |  | Threshold<br>(mg/kg)<sup>E</sup> | OAV |  |  |
| --- | --- | --- | --- | --- | --- | --- | --- | --- | --- | --- | --- | --- | --- | --- |
|  |  | Innowax | DB-5 |  | Concen<sup>C</sup> | SD<sup>D</sup> | Concen | SD | Concen | SD |  | Y1 | Y2 | Y3 |
| 1 | Butanal | 832 | 596 | MS, RI, Std | 0.065 | 0.006 | 0.016 | 0.001 | - | - | 0.017 | 4 | <1 | - |
| 2 | Ethyl acetate | 907 | 628 | MS, RI, Std | 3.209a<sup>F</sup> | 0.385 | 1.547b | 0.124 | 0.218c | 0.02 | 3.3 | <1 | <1 | <1 |
| 3 | Pentanal | 935 | 732 | MS, RI, Std | 0.014 | 0.001 | - | - | - | - | 0.012 | 1 | - | - |
| 4 | Ethyl propanoate | 951 | 713 | MS, RI, Std | 0.253b | 0.02 | 0.226b | 0.02 | 1.661a | 0.094 | 0.045 | 6 | 5 | 37 |
| 5 | Ethyl isobutanoate | 956 | 760 | MS, RI, Std | 0.085b | 0.009 | 0.349ab | 0.046 | 0.498a | 0.063 | 0.025 | 3 | 14 | 20 |
| 6 | Propyl acetate | 957 | 650 | MS, RI, Std | 0.545b | 0.038 | 0.074c | 0.008 | 1.683a | 0.112 | 2 | <1 | <1 | <1 |
| 7 | Methyl butanoate | 990 | 723 | MS, RI, Std | - | - | 0.011 | 0.001 | 0.008 | 0.001 | 0.001 | - | 11 | 8 |
| 8 | Methyl 2-methylbutanoate | 1005 | 739 | MS, RI, Std | 0.134a | 0.016 | 0.163a | 0.014 | 0.112a | 0.008 | 0.004 | 34 | 41 | 28 |
| 9 | Isobutyl acetate | 1015 | 776 | MS, RI, Std | - | - | 0.186 | 0.015 | - | - | NF | - | - | - |
| 10 | Butanol | 1024 | 676 | MS, RI, Std | 2.668b | 0.23 | 0.643c | 0.05 | 5.247a | 0.504 | 24 | <1 | <1 | <1 |
| 11 | Ethyl butanoate | 1028 | 804 | MS, RI, Std | 5.512ab | 0.605 | 6.906a | 0.564 | 1.914b | 0.181 | 0.4 | 14 | 17 | 5 |
| 12 | (E)-2-Butenal | 1031 | 624 | MS, RI, Std | 0.012 | 0.001 | - | - | - | - | 0.525 | <1 | - | - |
| 13 | 2,3-Pentanedione | 1041 | 698 | MS, RI, Std | 0.002 | 0 | - | - | - | - | 0.03 | <1 | - | - |
| 14 | Ethyl 2-methylbutanoate | 1058 | 843 | MS, RI, Std | 2.831b | 0.243 | 6.867a | 0.589 | 1.419c | 0.107 | 0.0003 | 94 | 229 | 47 |
| 15 | Butyl acetate | 1075 | 816 | MS, RI, Std | 1.932ab | 0.155 | 3.135a | 0.274 | 1.053b | 0.087 | 0.1 | 19 | 31 | 11 |
| 16 | Hexanal | 1078 | 803 | MS, RI, Std | 3.950a | 0.356 | 4.494a | 0.496 | 1.397b | 0.159 | 0.479 | 8 | 9 | 3 |
| 17 | 2-Methylpropanol | 1099 | 647 | MS, RI, Std | 0.184 | 0.02 | - | - | 0.523 | 0.044 | 16 | <1 | - | <1 |
| 18 | (E)-2-Pentenal | 1131 | 754 | MS, RI, Std | 0.014 | 0.002 | - | - | - | - | 1.5 | <1 | - | - |
| 19 | Ethyl pentanoate | 1133 | 903 | MS, RI, Std | 0.209 | 0.017 | - | - | - | - | 0.0087 | 24 | - | - |
| 20 | Butyl propanoate | 1138 | 908 | MS, RI, Std | 1.339a | 0.121 | 1.288a | 0.148 | 0.352b | 0.042 | 0.025 | 54 | 52 | 14 |
| 21 | (E)-2-Hexenal | 1192 | 844 | MS, RI, Std | 9.804a | 0.882 | 2.050b | 0.139 | 2.311b | 0.17 | 0.082 | 120 | 25 | 28 |
| 22 | 2-Methylbutanol | 1208 | 668 | MS, RI, Std | 2.301b | 0.207 | 3.787b | 0.53 | 13.421a | 1.136 | 32.5 | <1 | <1 | <1 |
| 23 | Ethyl hexanoate | 1220 | 1002 | MS, RI, Std | 4.878b | 0.527 | 7.016a | 0.926 | 0.859c | 0.069 | 0.22 | 22 | 32 | 4 |
| 24 | Butyl 2-methylbutanoate | 1220 | 1043 | MS, RI, Std | 0.894ab | 0.064 | 3.527a | 0.302 | 0.137b | 0.012 | 0.017 | 53 | 208 | 8 |
| 25 | Butyl butanoate | 1224 | 990 | MS, RI, Std | 2.037a | 0.247 | 2.632a | 0.204 | 0.319b | 0.029 | 0.11 | 19 | 24 | 3 |
| 26 | (E)-2-Heptenal | 1243 | 957 | MS, RI, Std | 0.085 | 0.009 | - | - | - | - | 0.051 | 2 | - | - |
| 27 | Heptanol | 1273 | 925 | MS, RI, Std | 0.655 | 0.079 | 0.269 | 0.027 | - | - | 0.33 | 2 | <1 | - |
| 28 | Hexanol | 1360 | 851 | MS, RI, Std | 1.22 | 0.095 | - | - | 6.655 | 0.509 | 2.5 | <1 | - | 3 |
| 29 | (E)-2-Hexen-1-ol | 1377 | 853 | MS, RI, Std | - | - | 2.121 | 0.208 | 3.377 | 0.308 | NF | - | - | - |
| 30 | Nonanal | 1387 | 1106 | MS, RI, Std | 0.203a | 0.017 | 0.189a | 0.017 | 0.004b | 0 | 0.04 | 5 | 5 | <1 |
| 31 | Octanol | 1388 | 981 | MS, RI, Std | 1.105 | 0.105 | 1.147 | 0.082 | - | - | 0.11 | 10 | 10 | - |
| 32 | (Z)-3-Hexenol | 1391 | 858 | MS, RI, Std | 0.172 | 0.021 | - | - | - | - | 0.07 | 2 | - | - |
| 33 | Ethyl octanoate | 1445 | 1197 | MS, RI, Std | 0.011a | 0.001 | 0.013a | 0.002 | 0.004b | 0 | 0.015 | <1 | <1 | <1 |
| 34 | Acetic acid | 1450 | 600 | MS, RI, Std | 34.181 | 2.016 | 37.294 | 3.035 | 48.76 | 3.543 | 26 | 1 | 1 | 2 |
| 35 | Benzaldehyde | 1498 | 962 | MS, RI, Std | 18.508a | 2.591 | 6.568b | 0.631 | 8.492b | 0.934 | 3.5 | 5 | 2 | 2 |
| 36 | Nonanol | 1502 | 1154 | MS, RI, Std | 0.655 | 0.05 | 0.951 | 0.089 | - | - | 1 | <1 | <1 | - |
| 37 | Benzyl acetate | 1510 | 1162 | MS, RI, Std | 0.367b | 0.028 | 1.037a | 0.086 | 0.324 | 0.001 | 0.03 | 12 | 35 | 10 |
| 38 | Ethyl decanoate | 1636 | 1398 | MS, RI, Std | 0.017a | 0.002 | 0.015a | 0.001 | 0.001b | 0 | 0.023 | <1 | <1 | <1 |
| 39 | α-Farnesene | 1763 | 1508 | MS, RI, Std | 3.234 | 0.291 | 2.783 | 0.242 | 1.892 | 0.148 | - | - | - | - |
| 40 | Decanol | 1765 | 1263 | MS, RI, Std | - | - | 0.116 | 0.014 | - | - | 0.775 | - | <1 | - |
| 41 | β-Damascenone | 1815 | 1387 | MS, RI, Std | 0.071a | 0.008 | 0.043b | 0.004 | 0.037b | 0.003 | 0.01 | 7 | 4 | 4 |

A: Retention indices of compounds on DB-5 and Innowax Column. B: MS, mass spectrum; RI, retention index; Std, authentic standards. C: The concentration of compounds in the delicious apple juices. D: Standard deviation. E: Van Gemert, L. J. Complilations of Odor Threshold Values in Air, Water and Other Media; Van SettenKwadraat: Houten, The Netherlands, 2003. F: The lowercase letters (a-c) following the values in the three delicious apple juices entries in the same row are significantly different according to the Duncan test (p < 0.05)

Dimethyl sulfide (DMS) was a volatile sulfur-containing compound, which was widely found in foods (Fatima et al., 2010; Mestres et al., 2000). DMS was considered as “asparagus, corn and molasses” aroma, which was beneficial to aroma profile of food at low concentrations. DMS could be chemically generated from a variety of organic sulfur precursors, such as S-methylmethionine. Furthermore, DMS assumed to convent from methionine in storage and transport forms (Segurel et al., 2004). Amongst the delicious apple juices, the contents of DMS ranged from 2.73 μg kg^−1^ in sample Y2 to 3.95 μg kg^−1^ in sample Y3. Although the contents of DMS was of trace level, the OAVs of this compound in three samples were all higher than one. The results indicated that DMS greatly contribute to the aroma of delicious apple juices.

Methional was sulfur-containing volatile compound widely found in red wine and grapefruit (Fatima et al., 2010; Mestres et al., 2000). In GC-O analysis, methional described as a typical “vegetable, cooked potato” aroma. At low concentration, methional considered to be a compound beneficial to the overall aroma. Methionol, as a reduction product of methional, brought the aroma of “vegetable and cabbage” aroma to delicious apple juices. According to previous studies, methionol could be obtained by decarboxylation of 4-methylthio-2-oxobutyric acid (MOBA) by alcohol dehydrogenase. Alternatively, methional could be oxidized to acid and reduced to alcohol according to the redox potential in cells or media (Landaud et al., 2008). From Table 3, the concentrations of methional were 0.84, 1.05 and 2.34μg kg^−1^ in three samples, while methionol were 87.4, 38.2 and 29.4 μg kg^−1^. Correspondingly, the OAVs of methional were 4.2, 5.3, and 11.7, respectively. Thus, methional considered as a great contributor to aroma of delicious apple juices. Compared with methional, methionol contributed little to the aroma of delicious apple juices, because OAVs of methionol were all less than 1.

2-(Methylthio)ethanol (MTE) was a thioether alcohol with the characteristic aroma of “vegetable and tomato”, which had been identified in different melon varieties (Ye et al., 2016). The highest concentration of MTE (27.89 g kg-1) was found in the Y3 sample, while the lowest concentration (16.85 g kg-1) was found in the Y2 sample. Correspondingly, the OAVs of MTE ranged from 1.2 to 1.9. Thus, MTE might potentially contribute to the delicious apple juices.

#### 3.3.2 Quantitative analysis and OAV of non-sulfur compounds

The concentrations and OAVs for non-sulfur compounds were listed in Table 4. The main compounds in delicious apple juice were esters, alcohols, aldehydes. Among those compounds, benzaldehyde (6.568-18.508 mg/kg), 2-methylbutanol (2.301-13.421 mg/kg), butanol (0.643-5.247 mg/kg), *(E)*-2-hexenal (2.050-9.804 mg/kg), ethyl butanoate (1.914-6.906 mg/kg), ethyl 2-methylbutanoate (1.419-6.867 mg/kg) presented relatively higher amounts than other compounds. The results were agreed with other studies which found that esters, alcohols, aldehydes, ketones compounds could be considered as the main compounds to delicious apple (Espinodíaz et al., 2016). The OAV was considered as useful method to evaluate the contribution of aroma compound to whole aroma. According to the previous literature (Guth, 1997), compounds were considered as contributors to aroma with OAVs above 1. From the Table 4, compounds ethyl 2-methylbutanoate (47-229), butyl 2-methylbutanoate (8-208), (E)-2-hexenal (25-120), butyl propanoate (14-54), methyl 2-methylbutanoate (28-41), ethyl hexanoate (4-32), ethyl butanoate (5-17) showed high OAVs in three delicious apple juices.

**Table 4.** The concentration (mg/kg), standard deviation and OAV of sulfur compounds detected in three delicious apple juices by FPD.

| No | Compounds | RI |  | Concentration (µg/kg) |  |  |  |  |  | Threshold<br>(µg/kg) | OAV |  |  |
| --- | --- | --- | --- | --- | --- | --- | --- | --- | --- | --- | --- | --- | --- |
|  |  | Wax | DB-5 | Y1 | SD | Y2 | SD | Y3 | SD |  | Y1 | Y2 | Y3 |
| 1 | Methanethiol | 696 | 502 | 6.36 | 0.53 | 6.29 | 0.63 | 4.55 | 0.14 | 4 | 1.6 | 1.6 | 1.1 |
| 2 | Dimethyl sulfide | 716 | 521 | 3.28 | 0.30 | 2.73 | 0.15 | 3.95 | 0.44 | 1.1 | 3.0 | 2.5 | 3.6 |
| 3 | Ethanethiol | 719 | 510 | 0.32 | 0.05 | 0.60 | 0.03 | 0.43 | 0.03 | 2 | 0.2 | 0.3 | 0.2 |
| 4 | Propanethiol | 863 | 616 | 0.54 | 0.07 | 2.65 | 0.17 | 1.04 | 0.28 | 3.1 | 0.2 | 0.9 | 0.3 |
| 5 | 3-Ethylthiophene | 1212 | 911 | - | - | 8.65 | 0.67 | 4.32 | 0.50 | - | - | - | - |
| 6 | Methional | 1444 | 905 | 0.84 | 0.13 | 1.05 | 0.01 | 2.34 | 0.08 | 0.2 | 4.2 | 5.3 | 11.7 |
| 7 | 2-(Methylthio)ethanol | 1513 | 818 | 143.20 | 11.60 | 232.10 | 11.30 | 43.90 | 3.84 | 120 | 1.2 | 1.9 | 0.4 |
| 8 | Methionol | 1721 | 985 | 87.40 | 6.43 | 38.20 | 2.43 | 29.40 | 2.56 | 500 | 0.2 | 0.1 | 0.1 |
| 9 | 3-Methyl-2-butene-1-thiol | 1904 | 962 | 13.20 | 0.12 | 8.43 | 0.12 | 5.43 | 0.13 | - | - | - | - |

The OAVs of methyl 2-methylbutanoate, ethyl butanoate, ethyl 2-methylbutanoate, butyl acetate, ethyl hexanoate, butyl 2-methylbutanoate and butyl butanoate in Y2 delicious apple juice were significantly higher than those in other samples. According to the previous research, ester compounds widely distributed in fruits, which were responsible for fruity, fresh aroma to fruits. Thus, those compounds were important compounds for fruit aroma, which consisted of the fundmental aroma of delicious apples.

Besides ester compounds, the OAVs of (E)-2-hexenal, octanol, hexanal, nonanal, (E)-2-heptenal were significantly higher than those of the other compounds. According to previous studies, many of fruits contain a large number of aldehydes, which contributed greatly to the formation of aroma (Hanoglu & Pucarelli., 2007).

Generally speaking, aldehydes containing 6 to 10 carbon have “green, fat or tallow” odors. Although most aldehydes produce those special and distinctive odors at low concentrations, they may also lead to rancidity, paint or other unpleasant odors at high concentrations due to their low thresholds (Zhu et al., 2015).

One important terpenoid, i.e., β-damascenone was detected in all three samples. Although the amounts of β-damascenone were low, the OAVs of this compounds were high due to the extreamly threholds (0.01 mg/kg). The highest OAV of this compound was obtained in the sample Y1 (OAV: 7), and lowest one in the sample Y1 and Y3 (OAV: 4). Thus, β-damascenone contributed greatly to the aroma of delicious apple samples.

### 3.4 Sensory analysis

Sensory evaluation was performed by organoleptic assessments of quality of three kinds of delicious apple juices using seven descriptors, including “wax”, “fruity”, “sour”, “green”, “sweet”, “sulfur” and “floral”. ANOVA was employed to ensure reliability of sensory data. As seen from Table 5, the significant intensity differences (p<0.01) existed in all descriptors. However, panellists presented no significant difference in all sensory descriptor. Through the interaction between the panel and the samples, the consistency of the scores of different samples was judged (Bárcenas et al., 2000). From the Table 5, no significant interaction between panellist×samples was also found. Thus, the sensory data was valid.

**Table 5.** Analysis of variance of the main effects and their interactions for sensory descriptor in descriptive analysis.

| Sensory descriptor | F values |  |  |
| --- | --- | --- | --- |
|  | Sample(S) | Panellist(P) | S×P |
| wax | 8.4 | 0.37 | 2.03 |
| sweet | 6.3 | 0.56 | 1.93 |
| fruit | 34.9* | 0.54 | 3.97 |
| floral | 59.2* | 0.62 | 4.76 |
| green | 69.8* | 0.68 | 2.63 |
| sour | 8.1 | 0.74 | 1.07 |
| sulfur | 3.5 | 0.95 | 2.09 |

\* Significant at $p < 0.05$ .

The intensities of “wax”, “sweet” and “green” in Y1 sample were greatly higher than other two samples. The compounds closely related to “wax” descriptor were 2-methylpropanol, heptanol, hexanol and 2-methylbutanol. As described in GC-O, β-damascenone was the most important contributor to “sweet” descriptor in delicious apple juices. The conclusion was consistent with other research which demonstrated that β-damascenone considered as “sweet” descriptor in jujub (Zhu & Xiao., 2018). Similarly, (E)-2-hexenal and hexanal contributed greatly to “green” note in delicious apple juices. The Y2 sample was accompanied by the “fruity” and “floral” descriptors. As described in GC-O, ester compounds were responsible for “fruity” descriptor, which was fundamental and important aroma to delicious apple juices. Ester compounds, such as ethyl 2-methylbutanoate, methyl 2-methylbutanoate, butyl propanoate and ethyl butanoate, exhibited a remarkably high correlation to “fruity” descriptor. According to result of GC-O, the compounds closely correlated to “floral” descriptor were benzyl acetate and unknown1. According to previous investigations, the “sour” descriptor related to acid compounds, which included acetic acid in delicious apple juices. In addition, methional, ethanethiol, methanethiol, dimethyl sulfide, 2-(methylthio)ethanol and methionol might contribute significantly to the “sulfur” descriptor in delicious apple juices. According the result of GC-O and OAV, methional and dimethyl sulfide were the most important contributors to “sulfur” descriptor.

### 3.5 Threshold variation of compounds added to reconstitution solution

From the above analysis, “fruity” descriptor was fundamental and important aroma to delicious apple juices. Esters were the characteristic compounds of fruit aroma. Thus, four volatiles (ethyl 2-methylbutanoate, ethyl hexanoate, ethyl butanoate and ethyl octanoate) with supra-threshold and sub-threshold concentrations were selected to evaluate their contributions. According to the previous research, compounds with low concentrations (sub-threshold) might also be contributor to the solution aroma (Lytra et al., 2013).

By doing so, the authors attempted to describe the contributions of single compounds to the entire aroma intensity more intuitively. As shown in Figure 1, all of the four aroma compounds presented dissimilar degrees of influence on the solution before and after their addition, with ratios of 5.89, 1.58, 3.16 and 1.77, respectively. It was clear that ethyl 2-methylbutanoate most significantly decreased the threshold of the solution, that is, it showed the most apparent enhancing effect on the aroma intensity of reconstitution solution. This was possibly because ethyl 2-methyl butanoate had a large OAV or it generated some synergistic effect with other compounds, so that aroma intensity of reconstitution solution was enhanced while the whole threshold of the solution dropped. In comparison, the threshold of the solution exhibited the least reduction after the addition of ethyl hexanoate and ethyl butanoate. This could possibly explained by their low OAVs, or because they underwent, or generated, little interaction with other compounds in the solution.

**Figure 1.**
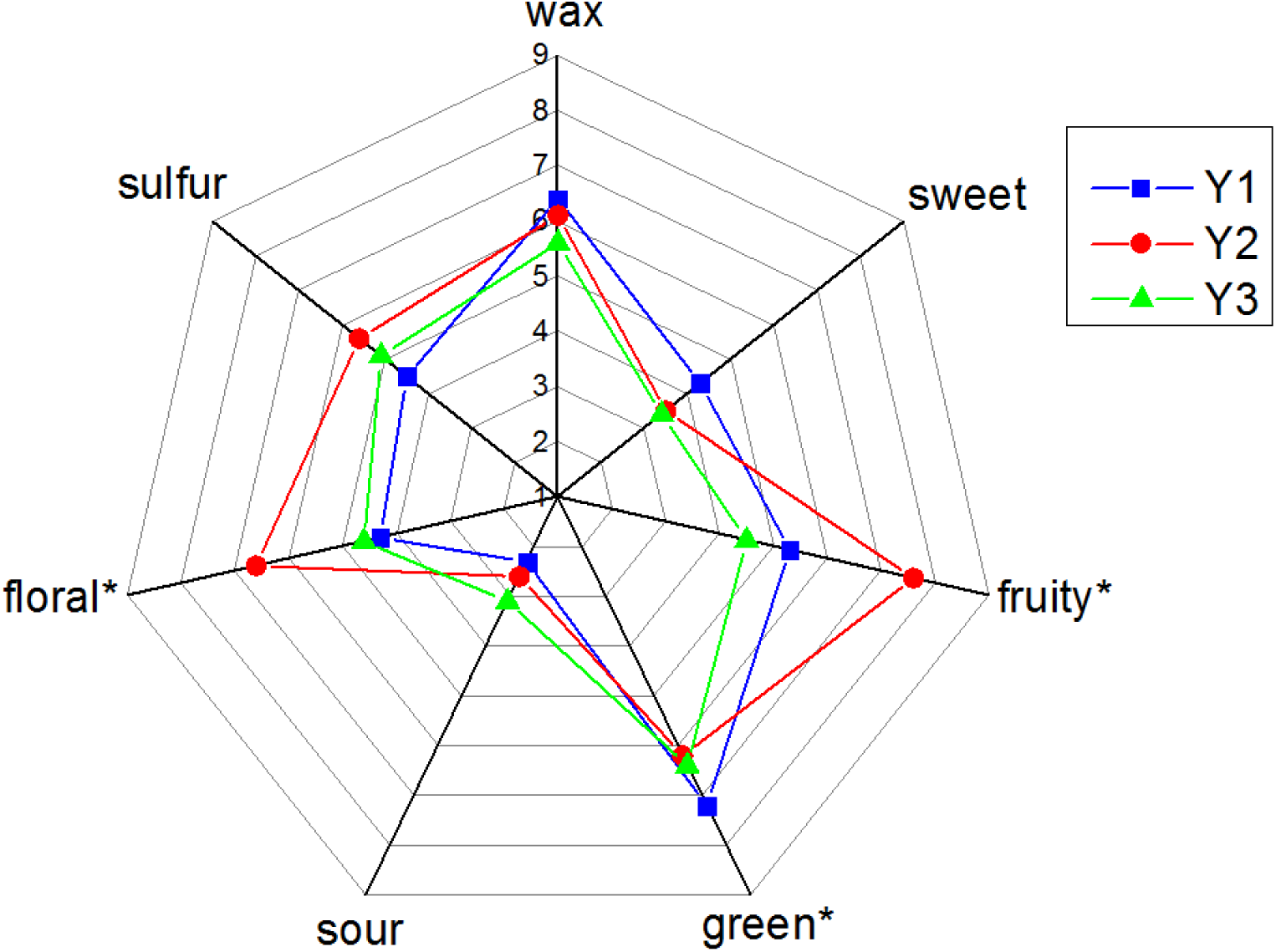
The aroma profiles of three delicious apple juices obtained from the Y1, Y2 and Y3 samples. The sensorial parameters indicated with an (*) are significantly different among some trials is verified for *p* < *0.05*.

**Figure 2.**
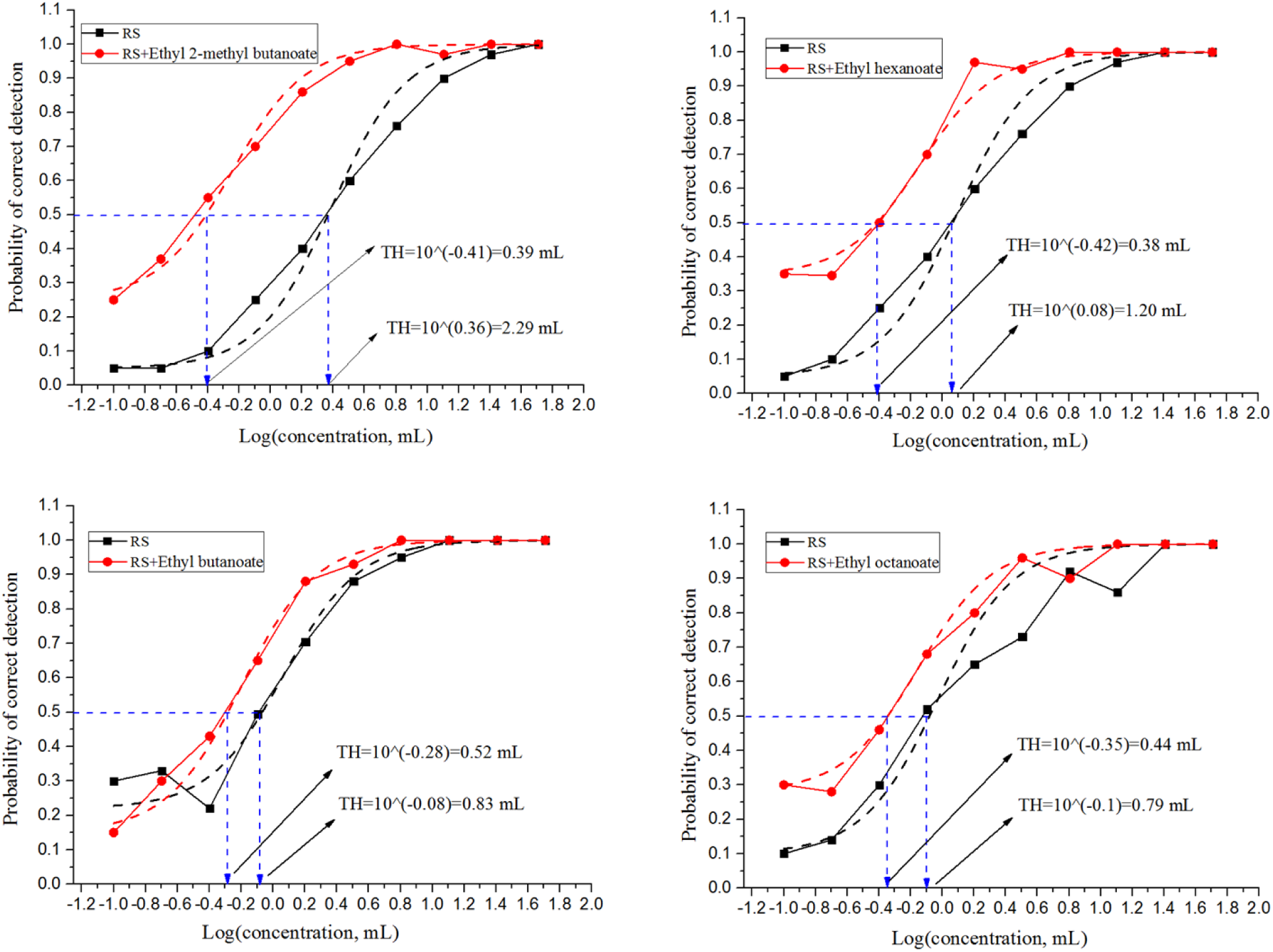
The comparing of detection probability of reconstitution solution before and after adding ethyl 2-methyl butanoate, ethyl butanoate, ethyl hexanoate and ethyl octanoate.

In particular, although the addition concentration of ethyl octanoate was lower than its threshold (OAV < 1), it decreased the threshold of the solution by 1.77. This suggested that it probably interacted with other compounds in reconstitution solution which reduced the threshold of the solution. Therefore it also made important contribution to aroma of solution. This coincided with results of previous research, which studied the blending of 12 esters found in wines on the impact of fruity aroma and concluded that ethyl propanoate with sub-threshold concentration, significantly strengthened fruity aromas and decreased the overall threshold of wines (Lytra et al., 2013).

## 4. Conclusion

GC-MS and GC-FPD were employed to identify the volatile compounds in delicious apple juices. The results indicated that 45, 41 and 37 volatile compounds were determined in the three delicious apple juices, respectively. Amongst those compounds, benzaldehyde, 2-methylbutanol, butanol, *(E)*-2-hexenal, ethyl butanoate, ethyl 2-methylbutanoate presented relatively higher amounts than other compounds. In addition, GC-O and OAV were used to determine the aroma-active compounds in fruit, suggesting that ethyl 2-methylbutanoate, butyl 2-methylbutanoate, (E)-2-hexenal, butyl propanoate, methyl 2-methylbutanoate and methional showed high OAVs and AIs in three delicious apple juices. Thus, those compounds contributed significantly to the aroma of delicious apple juices.

## Acknowledgements

The investigation was funded by Key Scientific Research Project of China Tobacco Yunnan Industrial Co., Ltd. [2018xy03, 2018JC04].

